# Genome assembly of *Eucalyptus recurva* provides insights into inbreeding and conservation priorities in Australia’s rarest *Eucalyptus*

**DOI:** 10.64898/2026.09.21.753279

**Authors:** Eilish S. McMaster, Ashley Jones

## Abstract

*Eucalyptus recurva* (Mongarlowe Mallee) is Critically Endangered, with only six known adult individuals persisting across two sites in the Southern Tablelands of New South Wales, Australia. Its extreme rarity, uniquely long lifespan, and limited reproductive output make it a priority for conservation genomics. Here we report the first genome assembly of *E. recurva*: a haplotype-resolved, gapless, telomere-to-telomere (T2T) assembly produced from Oxford Nanopore Technologies (ONT) long-read sequencing. Both haplotypes span 11 chromosomes (consistent with the conserved *Eucalyptus* karyotype of 2n = 22), with assembly sizes of 521.1 Mb (Hap 1) and 501.5 Mb (Hap 2), BUSCO completeness >99.6%, and quality values of >QV 62. Comparative analyses place *E. recurva* within section Maidenaria as sister to *Eucalyptus viminalis* and reveal high synteny between the two species. Inter-haplotype comparison identified 3.6 million SNPs and modest structural variation, suggesting that, despite extreme demographic bottlenecking, *E. recurva* retains meaningful genomic heterozygosity. We also characterise the chloroplast genome, which exhibits heteroplasmy, and report an apparently bipartite mitochondrial genome. This reference genome provides an essential resource for conservation management, population genetics, and the study of eucalypt genome evolution.

## Introduction

*Eucalyptus recurva*, commonly known as the Mongarlowe Mallee, is one of Australia’s most imperilled plant species. With only six known adult individuals remaining across two sites in the Southern Tablelands of New South Wales (Mongarlowe and Windellama), it holds the distinction of being Australia’s rarest eucalypt (Crisp, 1988; Fensham, 2019). Listed as Critically Endangered under both the Australian Environment Protection and Biodiversity Conservation Act 1999 and the IUCN Red List, *E. recurva* is characterised by strongly recurved leaves, smooth bark, and a multi-stemmed habit arising from an extensive lignotuber (Crisp, 1988). Lignotubers extending up to 12 m across suggest that surviving individuals may be hundreds to potentially thousands of years old (NSW National Parks & Wildlife Service, 2003). This extraordinary longevity, combined with limited reproductive success (Martyn Yenson et al., 2024; NSW National Parks & Wildlife Service, 2003), suggests that *E. recurva* represents an evolutionary lineage maintained in recent times primarily through vegetative persistence; an ‘ice age gum’ that may be a relic of past conditions (Nicolle and Jones, 2018).

The species is placed within *Eucalyptus* subgenus Symphyomyrtus, section Maidenaria (Crisp, 1988; Nicolle, 2021; Thornhill et al., 2019). It occupies a dry, low-rainfall environment near Braidwood on the Southern Tablelands (Benson, 2024). The known individuals grow in highly restricted, geographically fragmented patches, with no confirmed recent natural recruitment. Multiple threats compound the conservation challenge: projected declines in habitat suitability under future climate scenarios (Archibald et al., 2024), increasing frequency and severity of bushfires (Godfree et al., 2021), and increasing spread of fungal pathogens such as *Phytophthora cinnamomi* (causing dieback; (McDougall et al., 2024)) and *Austropuccinia psidii* (myrtle rust; (Soewarto et al., 2025)). Understanding the genetic diversity and adaptive potential of the surviving individuals is therefore urgent.

Conservation genomics is transforming the management of Critically Endangered species in Australia, providing cost-effective insights into genetic health and evolutionary potential (Doyle et al., 2025). Advances in long-read sequencing have enhanced this further, enabling haplotype-resolved, reference-quality genome assemblies that reveal structural variation invisible to short-read approaches (Leitwein et al., 2020). Haplotype information provides a richer view of population dynamics than individual SNPs alone by capturing patterns of gene flow, inbreeding, and adaptation (Chen et al., 2025, 2023). In *Eucalyptus*, telomere-to-telomere (T2T) assemblies are now available for *E. regnans* and *E. viminalis*, revealing extensive structural variation between haplotypes and structural rearrangements as drivers of genome evolution, despite a conserved karyotype of 2n = 22 across the genus (Ferguson et al., 2024b; Zhuang et al., 2026).

The conservation genomics of *E. recurva* was first comprehensively addressed using DArTseq reduced-representation sequencing to all six wild adults alongside a range of ex situ individuals including recent seedling cohorts, cultivated hybrids, and grafted material (McMaster et al., 2026). This study confirmed that all six wild adults are genetically distinct genotypes, resolving earlier uncertainty about the clonal status of the Windellama individuals, and revealed population structure between the Mongarlowe and Windellama sites. Parentage analysis of ex situ seedling cohorts showed natural hybridisation with sympatric congeners *E. rubida* subsp. *rubida* and *E. mannifera* subsp. *mannifera*. A broader phylogenetic network identified *E. sturgissiana* as the closest known relative of *E. recurva*. Together, these findings established that *E. recurva* faces compounding pressures of inbreeding, constrained genetic diversity, and reproductive failure consistent with early-stage extinction vortex dynamics, and that managed introgression represents a lower-risk conservation option than continued inaction (McMaster et al., 2026).

For *E. recurva* specifically, no genome sequence, karyotype confirmation, or characterisation of genome architecture has previously been reported. Whole-genome sequencing extends beyond SNP-based approaches by enabling the detection of structural variants and chromosomal rearrangements that are largely invisible to reduced-representation methods; the phasing of haplotypic diversity across full chromosomes; the assembly and analysis of organelle genomes; the characterisation of gene content including disease resistance loci; and the creation of a reference resource to support future population resequencing and functional genomic study. Here we present the first genome assembly of *E. recurva*: a haplotype-resolved, gapless T2T assembly generated from a single individual using Oxford Nanopore Technologies (ONT) long-read sequencing. We characterise genome quality, investigate haplotypic variation, determine ploidy, assemble organelle genomes, reconstruct the phylogenetic position of *E. recurva* within a broad *Eucalyptus* genome dataset, and perform synteny analyses comparing *E. recurva* to closely and distantly related species. Together, these analyses provide foundational genomic resources to support the long-term conservation of this Critically Endangered species.

## Results and Discussion

### High-coverage sequencing and genome profiling

High-molecular-weight DNA was extracted from leaf tissue of a single *E. recurva* individual (Fig S1), which was size selected for ≥40 kb fragments and sequenced on the ONT PromethION P2 Solo platform using two R10.4.1 flow cells. Total sequencing output was 217 Gbp from 8.28 M reads (Table 1). After stringent filtering for the longest and most accurate reads (≥40 kb and ≥Q20), 117.3 Gbp from 2.14 M reads was retained for genome assembly (N50 54,079 bp, median read quality Q25), corresponding to approximately 225 × total coverage (113 × per haplotype). A further subset of ultra-long reads (≥100 kb, ≥Q7; 29,789 reads, 3.50 Gbp) was retained for gap closure. The longest read in the ≥40 kb Q20 subset was 306,218 bp, and in the ultra-long subset it was 663,814 bp. This high level of coverage, read length, and quality is consistent with recent studies that achieve complete, gapless T2T plant genome assemblies (Ma et al., 2026).

**Table 1.** Oxford Nanopore Technologies sequencing metrics for *Eucalyptus recurva*.

| Metric | ONT (all) | ONT $\geq 40$ kb Q20 | ONT $\geq 100$ kb Q7 |
| --- | --- | --- | --- |
| Reads | 8,281,379 | 2,143,959 | 29,789 |
| Yield (Gbp) | 217.4 | 117.3 | 3.50 |
| Mean read length (bp) | 26,247 | 54,724 | 117,573 |
| Median read length (bp) | 22,132 | 51,214 | 110,182 |
| N50 (bp) | 46,191 | 54,079 | 112,185 |
| Mean read quality | Q12.8 | Q23.8 | Q18.2 |
| Median read quality | Q23.6 | Q25.0 | Q22.1 |
| Longest read (bp) | 2,058,801 | 306,218 | 663,814 |
| Estimated coverage (521 Mbp) | 417 × | 225 × | 6.72 × |

We performed a k-mer frequency spectrum analysis of the sequencing reads with GenomeScope2 (Fig. S2) to estimate genome size, heterozygosity, and ploidy, which was further validated by k-mer pair ratio analysis using Smudgeplot (Fig. S3). Both analyses confirmed that *E. recurva* is diploid (2n = 22), with two distinct k-mer peaks consistent with a heterozygous diploid genome. Genome size was estimated at approximately 521 Mbp consistent with other *Eucalyptus* genomes (Ferguson et al., 2023, 2024b). Despite the extreme demographic bottleneck represented by only six surviving wild individuals, *E. recurva* retains substantial heterozygosity at 0.89% (HoA = 3,600,801 SNPs / 404,995,050 syntenic bp = 0.89%), consistent with patterns observed across the genus; even the most genetically depauperate eucalypts appear more heterozygous than most other plant species.

### Haplotype-resolved, gapless T2T assembly

*De novo* assembly with hifiasm (ONT) using the ≥40 kb Q20 read set produced two highly contiguous, haplotype-resolved assemblies (Table S1). The largest contigs were T2T chromosomes of the nuclear genome (10/11 for Haplotype 1, 9/11 for Haplotype 2). Dual scaffolding (--dual-scaf) resolved one break point in Haplotype 2 (relative to Haplotype 1). Final chromosome ordering was performed using RagTag, guided by the closely related *E. viminalis* T2T assembly (Zhuang et al., 2026) as a reference, with results confirmed using the more distantly related *E. regnans* T2T assembly (Ferguson et al., 2024a). This scaffolding approach resolved the final break point in both haplotypes, which appeared to be the centromeric region of chromosome 2. Both haplotypes were placed into 11 chromosomes, consistent with the conserved *Eucalyptus* karyotype of 2n = 22. The three residual gaps, at the likely chromosome 2 centromere for each haplotype and a gap in chromosome 4 for haplotype 2, were closed using TGS-GapCloser with ultra-long ONT reads ≥100 kb Q7 (Table S2). All 22 telomeres were detected for each haplotype (Figure S4), indicating a fully gapless, T2T nuclear genome. Genome assembly metrics are summarised in Table 2. The final Haplotype 1 was 521,095,720 bp (N50 52.3 Mbp), and Haplotype 2 was 501,541,970 bp (N50 50.3 Mbp). The size difference between haplotypes (~19.5 Mbp) is consistent with haplotypic structural variation observed in other *Eucalyptus* species (Ferguson et al., 2024b), and other wild trees Australian trees (Chen et al., 2023), grasses (Chen et al., 2026), and orchids (Z. Zhang et al., 2026).

**Table 2.** Final assembly metrics for the two haplotypes of *Eucalyptus recurva* (karyotype 2n = 22).

| Metric | Haplotype 1 | Haplotype 2 |
| --- | --- | --- |
| **Sequences (chromosomes)** | 11 | 11 |
| **Total size (bp)** | 521,095,720 | 501,541,970 |
| **N50 (bp)** | 52,276,560 | 50,321,341 |
| **Telomeres detected** | 22/22 | 22/22 |
| **Telomere-to-telomere chromosomes** | 11/11 | 11/11 |
| **Assembly gaps** | 0 | 0 |
| **BUSCO completeness (eudicotyledons_odb12)** | 99.61% | 99.65% |
| **Single copy** | 97.68% | 97.90% |
| **Duplicated** | 1.93% | 1.75% |
| **Fragmented** | 0.32% | 0.29% |
| **Missing** | 0.07% | 0.07% |
| **Quality Value (QV, raw / adjusted)** | 62.07 / 61.85 | 68.62 / 67.72 |

Assembly completeness was assessed with Compleasm v0.2.7 using the eudicotyledons_odb12 BUSCO dataset (2,805 genes). Both haplotypes exceeded 99.6% completeness (Hap 1: 99.61%; Hap 2: 99.65%), with single-copy rates of 97.68% and 97.90% respectively (Table 2). The assembly completeness rivals many of the available T2T plant genomes (Xie et al., 2024), including gapless T2T crop genomes (Ma et al., 2026). The absence of two BUSCO genes on each haplotype is likely as a true observation, given the *E. recurva* genome is gapless T2T. Mapping the reads back to the assembly (Figure S5), 99.95% and 99.96% of reads aligned to haplotypes 1 and 2, respectively, with relatively even coverage genome-wide and the lowest coverage restricted to highly repetitive regions such as suspected centromeres. Assembly accuracy was assessed using yak (k = 31), which yielded quality values of QV 62.07 / 61.85 (Hap 1; raw/coverage-adjusted) and QV 68.62 / 67.72 (Hap 2), indicating per-base assembly accuracies >99.9999%. These values exceed QV40, the benchmark associated with the Vertebrate Genomes Project (Rhie et al., 2020), and exceed human T2T genome standards (QV50; (Cheng et al., 2026)), which underscore the substantial accuracy improvements delivered by modern ONT chemistries (Ferguson et al., 2022). Overall, the *E. recurva* assembly is among the highest-quality plant genomes reported to date.

### Organelle genomes

The chloroplast assembly revealed heteroplasmy, with two structural variants of equal size (161,691 bp each), consistent with the characteristic inversion of the small single-copy region commonly observed in plastomes (Figure S6). Annotation using GeSeq identified the expected suite of chloroplast genes using all available *Eucalyptus* chloroplast references on NCBI.

The mitochondrial assembly resolved into two circular components of 313,337 bp and 185,047 bp, yielding a combined mitogenome size of 498,384 bp (Figure 1a). The limited number of available *Eucalyptus* mitochondrial genomes are reported as single circular chromosomes of approximately 400-479 kb (*E. camaldulensis*: 463 kb (Fukasawa et al., 2024); *E. grandis*: 479 kb (Pinard et al., 2019)). The bipartite structure observed here may reflect genuine multi-chromosomal mitochondrial organisation or could represent a structural variant stabilised in this highly reduced lineage. Ultra-long reads and new plant-specific organelle assembly tools used in our approach are critical for resolving complex mitogenome architectures that are otherwise collapsed into a single contig, as evidenced by the highly diverse organelle genome structures being revealed across many plant genera (Kozik et al., 2019; Xian et al., 2025; Zhou et al., 2025).

**Figure 1.**
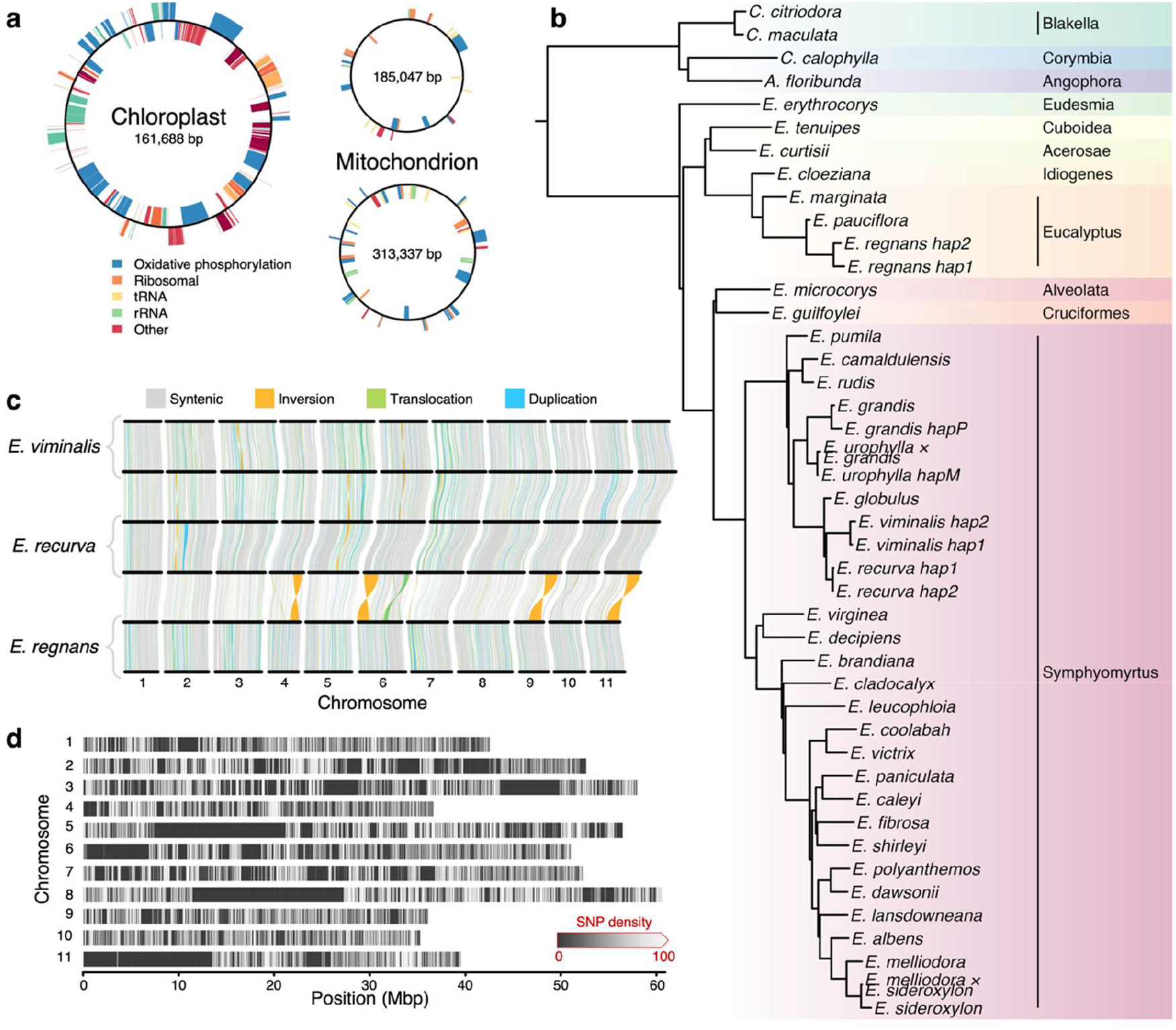
(a) Architecture and annotation of the *Eucalyptus recurva* organelle genomes: chloroplast (left) and mitochondrion (right; two circular parts). Genome schematics not to scale. (b) Species tree of *Eucalyptus* and closely related *Corymbia* and *Angophora* from single-copy orthologous genes, with subgenera highlighted. (c) Synteny and structural variation between the haplotypes of *E. viminalis, E. recurva*, and *E. regnans*. Whole-genome alignments were annotated with SyRI and filtered to intrachromosomal variants ≥20 kb at ≥90% sequence identity. (d) Genome-wide SNP density across the 11 chromosomes of *E. recurva*. Each row represents one chromosome, with shade indicating the number of SNPs per 10 kbp window (dark: low density; light: high density, ≥100 SNPs per 10 kb, equivalent to ≥1% heterozygosity). Extended dark tracts are consistent with runs of homozygosity.

Multipartite mitogenomes are not uncommon in plants and do not inherently impair fertility. However, should *E. recurva* prove to have reduced pollen viability specifically (rather than other forms of reproductive failure), mitochondrial rearrangements of the kind observed here could be implicated. Cytoplasmic male sterility (CMS) is a well-documented consequence of mitochondrial genome rearrangements in plants, with demonstrated effects on pollen viability across a range of taxa (Chen and Liu, 2014). The biological significance of the *E. recurva* mitogenome architecture therefore might warrant further investigation.

### Phylogenomic placement

*Eucalyptus recurva* was placed in a phylogenomic context using the publicly available 44 eucalypt genome assemblies spanning 38 species, including four at haplotype resolution and two interspecific hybrids, plus the outgroup *Syzygium grande*. All genomes had BUSCO completeness ≥92.8%. The resulting tree (Figure 1b) is highly concordant with established *Eucalyptus* taxonomy (Nicolle, 2021; Thornhill et al., 2019), with strong support for all major clades. *Eucalyptus recurva* resolves within section Maidenaria, positioned between *E. globulus* and *E. viminalis*, as expected from its morphological placement in that section. *E. sturgissiana* was identified as the closest known relative in the previous DArTseq study (McMaster et al., 2026), but the absence of a whole-genome assembly for that species means *E. viminalis* is the nearest sampled relative in the present dataset. Notably, the two haplotypes of *E. recurva* are far more similar to one another in terms of BUSCO gene sequence divergence than the haplotype pairs of *E. viminalis* and *E. regnans*, suggesting reduced inter-haplotype sequence divergence consistent with a recent or prolonged population bottleneck. Two *Corymbia* species resolved at the base of the tree, consistent with the *Blakella* grouping proposed by Crisp (Crisp et al., 2024).

### Synteny and inter-haplotype structural variation

Pairwise genome alignments were performed to investigate the synteny and structural variation between haplotypes within *E. recurva* and across two other T2T *Eucalyptus* genomes, the closely related *E. viminalis* (subgenus Symphyomyrtus, section Maidenaria) and the more distantly related *E. regnans* (subgenus Eucalyptus, section Eucalyptus; Figure 1c). Comparison of the two *E. recurva* haplotypes revealed them to be the most syntenic pair of the three species examined, with 74.67% of haplotype 1 and 79.07% of haplotype 2 annotated as syntenic (Figure S8, Table S3). The equivalent values were 72.55% and 74.62% in *E. regnans*, and 66.39% and 68.20% in *E. viminalis*. There were also fewer inversions in *E. recurva*, at 68 events totalling 5.32 Mbp, or 1.02% of haplotype 1, with translocations and duplications accounting for a further 5.14% and 9.20%. At the sequence level we identified 3,600,801 SNPs, 362,963 insertions, and 325,456 deletions between haplotypes, along with 3,111 highly diverged regions spanning 7.52 Mbp (Table S4). Fewer SNPs and less diverged sequence were recovered in *E. recurva* than in either *E. regnans* (4,843,471 SNPs; 9.83 Mbp) or *E. viminalis* (5,900,220 SNPs; 14.10 Mbp), which is consistent with the reduced haplotype divergence observed in the phylogenomic analysis. *E. recurva* nonetheless retains substantial heterozygosity, despite centuries to millennia of demographic isolation.

Synteny between species also followed a trend consistent with the phylogenomic analysis (Figure 1c). *E. recurva* and *E. viminalis* remained collinear across all 11 chromosomes, with 59.70-64.40% of each assembly syntenic and inversions limited to ~1-2% (Table S3). Compared to *E. regnans* (section Eucalyptus rather than Maidenaria), synteny became considerably fragmented: only 37.59-40.93% of *E. recurva* and 36.91-40.68% of *E. viminalis* remained syntenic, ~37-42% of each assembly could not be aligned, and highly diverged sequence rose from 15.80 Mbp to 114.39 Mbp (Table S4). Inversions increased to ~11% of the annotated length in both species (9.91-11.18% for *E. recurva*, 10.81-11.11% for *E. viminalis*), driven by four macro-scale inversions on chromosomes 4, 6, 9, and 11 (Figure 1c). Similarly, we observed a macro-scale translocation on chromosome 6, highlighting that chromosome 6 is the most structurally rearranged between the two sections Maidenaria and Eucalyptus. Chromosomal rearrangements of this size may offer new insights into differences in phenotypes between the species, for instance *E. regnans* being one of the tallest flowering trees on earth (Ferguson et al., 2024a; Williams et al., 2023), compared to the smaller stature of the other two species. However, macro-scale structural variants suppress recombination in heterozygotes (X. Zhang et al., 2026), and are likely to act as meiotic barriers to gene flow between *E. regnans* and *E. recurva*. The conserved synteny between *E. recurva* and *E. viminalis* suggests that hybridisation between these species is structurally feasible and is consistent with the previous finding of natural *E. recurva × E. rubida* and *E. recurva* × *E. mannifera* hybrids (McMaster et al., 2026), which are close relatives to *E. viminalis* (all series Viminales (Nicolle, 2021)). Therefore, regarding crossing or managed introgression, it is likely that *E. recurva* is restricted to only closely related species within section Maidenaria (McMaster et al., 2026), which limits the pool available for genetic rescue of an already severely bottlenecked species.

### Genome-wide SNP density reveals runs of homozygosity

In our synteny analyses, we observed several megabase-scale blocks within the *E. recurva* comparison were entirely syntenic, carrying no inversions, translocations, duplications, or SNPs. Nothing comparable was seen between the haplotypes of *E. viminalis* or *E. regnans*. We further investigated this lack of inter-haplotype variation, as they may be genuine or may reflect collapse of the two haplotypes during assembly. Reads (≥40 kb and ≥Q20) mapped to haplotype 1 were used to independently call SNPs, and counts were tallied in 10 kb windows for each chromosome (Figure S7) and summarised genome-wide (Figure 1d). SNP density varied substantially across all 11 chromosomes, with extended regions of near-zero density interspersed among windows of moderate-to-high heterozygosity. Large contiguous low-density tracts were most prominent on chromosomes 3, 5, 6, 8, and 11, spanning tens of megabases and consistent with runs of homozygosity (ROH). Boundaries of the largest tracts were manually inspected and catalogued (Table S5), revealing six regions spanning 3.51-15.83 Mbp in size, totalling 59.51 Mbp, or 11.42% of haplotype 1. Coverage was indeed high and relatively uniform throughout these regions (Figure S5), and both haplotypes assembled as complete, gapless chromosomes, indicating true homozygosity rather than errors in genome assembly.

Tracts of this length arise from haplotypes inherited identical by descent and are a signature of recent, close inbreeding. The ROH highlight where recessive deleterious alleles may occur and are therefore the most probable source of inbreeding depression in the species. Whether the limited reproductive output, strongly recurved leaves, and reduced mallee stature reflect variation fixed within these tracts remains to be tested. Of more immediate concern is the loss of allelic diversity at loci where heterozygosity is functionally important, particularly NLR-type resistance genes, which are allelically diverse in other Myrtaceae (Chen et al., 2023). Any resistance loci within these ROH will carry a single allele, which may compromise the immune response of *E. recurva* to fungal diseases such as *Austropuccinia psidii* (myrtle rust) and *Phytophthora cinnamomi*. Annotation of their gene content is a priority, and the ROH provide explicit targets against which genetic rescue, whether by crossing the remaining genotypes (McMaster et al., 2026) or managed introgression, can be directed and measured.

### Conservation implications

The *E. recurva* genome assembly presented here provides an important resource for conservation genomics of this Critically Endangered species. The six known wild individuals were recently characterised using unmapped DArTseq-derived SNPs, confirming that they are genetically distinct and providing initial insights into kinship structure and inbreeding dynamics (McMaster et al., 2026). This chromosome-scale, haplotype-resolved reference substantially expands that toolkit, enabling analyses from structural variant detection to demographic reconstruction that are not accessible from unmapped reduced-representation data.

The genomic heterozygosity observed in this individual is encouraging, as despite extreme rarity, *E. recurva* does not appear to have experienced complete loss of genetic variation at the sequence level. This parallels other rare Australian eucalypts, including *E. caesia* and *E. argutifolia*, which maintain moderate-to-high outcrossing rates independent of population size (Bezemer et al., 2016; Kennington and James, 1997). Notably, DArTseq analysis of the six known adults found that this individual had the lowest heterozygosity of the group (McMaster et al., 2026), raising the possibility that the remaining genotypes harbour greater diversity and fewer runs of homozygosity than reported here. Reference-guided resequencing of the remaining wild genotypes would allow this to be assessed population-wide and would map the runs of homozygosity identified here across all six individuals, targeting genetic rescue efforts by identifying which loci most urgently need novel variation through crossing among wild genotypes or managed introgression.

The high synteny with *E. viminalis* also raises the possibility of identifying genomic regions of functional importance shared across Maidenaria, informing which related species are best suited as introgression candidates. Characterising large structural variants between potential crossing partners adds a further layer of value: chromosomal inversions and translocations suppress recombination specifically in heterozygotes and can lock functionally important genes into non-recombining blocks (X. Zhang et al., 2026), acting as reproductive barriers. Screening candidate parents for such variants, as demonstrated here between *E. recurva* and *E. regnans*, would help anticipate reduced fertility or restricted recombination before a crossing program is attempted rather than after.

This reference also underpins rigorous genomic evaluation of any crossing program, enabling multi-generational monitoring of hybrid genomic composition, detection of structural incompatibilities between parental genomes, and assessment of outbreeding depression risk prior to field release. Conservation of genome-wide genetic variation remains the most robust approach to maintaining adaptive potential in threatened populations (Kardos et al., 2021), and high-quality reference genomes are increasingly central to translating that principle into management action.

## Conclusions

We present the first genome assembly of *E. recurva*, Australia’s rarest eucalypt: a haplotype-resolved, gapless, T2T reference of exceptional quality (BUSCO >99.6%, QV >62). The assembly resolves both haplotypes into 11 chromosomes, confirms a diploid karyotype (2n = 22), and reveals that the species retains meaningful genomic heterozygosity despite extreme demographic reduction, alongside runs of homozygosity that mark clear targets for future genetic rescue. Phylogenomic analysis confirms the placement of *E. recurva* in section Maidenaria as sister to *E. viminalis*, with high synteny between the two species. The bipartite mitochondrial genome and chloroplast heteroplasmy are notable features warranting further investigation. This reference genome represents a foundational resource for the genomics-informed conservation management of *E. recurva* and contributes to the growing compendium of high-quality Eucalyptus genomes enabling comparative and evolutionary studies across this ecologically important and taxonomically diverse genus.

## Materials and Methods

### Tissue collection

A scientific licence was obtained from the New South Wales Government, Department of Climate Change, Energy, the Environment and Water (licence number SL102956), in accordance with the Biodiversity Conservation Act 2016. Material was collected from a single *E. recurva* individual by a Senior Threatened Species Officer (G. Wright) from the Southern Tablelands region of New South Wales, Australia. An herbarium voucher for this individual is held in the National Herbarium of New South Wales (NSW 496534; collected by G. Leonard, 3 December 1999; described as Specimen 1 from the site, corresponding to individual “D” in McMaster et al. 2026) and is documented in the Australasian Virtual Herbarium. Leaf material was maintained cool and moist following collection and cryogenically stored at ™80°C at the Australian National University, Canberra.

### DNA extraction and sequencing

High-molecular-weight (HMW) DNA was extracted using a magnetic bead-based protocol (Jones et al., 2021). Briefly, leaf tissue was ground under liquid nitrogen, homogenate was washed with a sorbitol buffer, cells lysed with SDS buffer, proteins precipitated with potassium acetate, and DNA bound to magnetic beads and washed with ethanol prior to elution. Extracted HMW DNA was size selected for fragments ≥40 kb using a BluePippin (Sage Science) according to the manufacturer’s instructions. The HMW DNA was assessed by pulsed-field capillary electrophoresis (Femto Pulse, Agilent Technologies), before and after size selection.

An ONT native DNA sequencing library was constructed using the ‘Ligation sequencing DNA V14 (SQK-LSK114)’ protocol with the NEBNext Companion Module v2 (NEB E7672S). Sequencing was performed on an ONT PromethION P2 Solo (PRO-SEQ002) using two R10.4.1 PromethION flow cells (FLO-PRO114M). To maximise yield, each flow cell was washed with DNAse I and re-loaded with fresh library at least twice using the Flow Cell Wash Kit (EXP-WSH004), until the flow cell was expended.

### Base calling, quality control, and genome assembly

Base calling was performed with Dorado v7.6.8 (super-accuracy model v4.3.0, 400 bps). Raw reads were quality-filtered for ≥40 kb ≥Q20 for genome assembly, and ≥100 kb ≥Q7 for gap closure, using Chopper v0.8.0 (De Coster and Rademakers 2023). De novo haplotype-resolved genome assembly was performed with hifiasm (ONT mode) v0.25.0-r726 (Cheng et al., 2025) using the ≥40 kb ≥Q20 read set, with the *Eucalyptus* telomere sequence specified (--telo-m AAACCCT) and dual scaffolding enabled (--dual-scaf).

Final chromosome ordering was performed with RagTag v2.1.0 (Alonge et al., 2022), using the *E. viminalis* T2T genome (Zhuang et al., 2026) as the primary reference; scaffolding results were confirmed using the *E. regnans* T2T genome (Ferguson et al., 2024a). Residual gaps at the chromosome 2 centromere (one gap in Hap 1, two gaps in Hap 2) were closed with TGS-GapCloser v1.2.1 (Xu et al., 2020) using ONT reads filtered to ≥100 kb and ≥Q7.

### Assembly quality assessment

Assembly completeness was evaluated using Compleasm v0.2.7 (Huang and Li, 2023) with the eudicotyledons_odb12 BUSCO dataset ((Tegenfeldt et al., 2025); 2,805 genes). Assembly accuracy (quality value, QV) was estimated with yak v0.1 (k = 31; (Li, 2021)), using the ≥40 kb ≥Q20 read set. Ploidy and genome size were estimated from 21-mer frequency distributions using GenomeScope2 and SmudgePlot v0.3 (Ranallo-Benavidez et al., 2020). Genome assembly graphs were visualised with Bandage v0.8.1 (Wick et al., 2015).

### Organelle genome assembly and annotation

Chloroplast and mitochondrial genomes were assembled using TIPPo v1.3.0 (Xian et al., 2025). Organelle assemblies were annotated with GeSeq (Tillich et al., 2017). Chloroplast annotation used all available *Eucalyptus* chloroplast reference sequences on NCBI (*E. camaldulensis, E*. globulus, *E. grandis*) and the MPI-MP chloroplast land plants reference set, with the Chloë v0.1.0 tool for CDS/tRNA/rRNA support and tRNAscan-SE v2.0.7 for organellar tRNAs. Mitochondrial annotation used all malvid mitogenome references available on NCBI with default BLAT search settings.

### Phylogenomics

A phylogenomic dataset was compiled from 44 eucalypt genome assemblies spanning 38 species (including four haplotype-resolved assemblies and two interspecific hybrids (Ahrens et al., 2024; Driguez et al., 2021; Ferguson et al., 2024a, 2024b, 2024c, 2023; Healey et al., 2021; Lötter et al., 2023; Shen et al., 2023; Zhuang et al., 2026); all with BUSCO completeness ≥92.8%), together with the Myrtaceae outgroup *Syzygium grande* (Low et al., 2022). Single-copy orthologous genes were identified with BUSCO v5.8.2 (eudicotyledons_odb12, 2,805 genes). Protein sequences present in ≥97% of genomes were aligned with MUSCLE5 v5.1 (Edgar, 2022) and poorly aligned regions removed with trimAl v1.4.rev15 (Capella-Gutiérrez et al., 2009). Maximum likelihood gene trees were inferred with IQ-TREE v2.3.6 (Minh et al., 2020) and the species tree estimated under the multi-species coalescent model with ASTRAL-IV v1.23.4.6 (Zhang et al., 2025). The species tree was rooted on *S. grande* and visualised with ggtree v4.0.5.

### Synteny and structural variation analysis

Pairwise genome alignments were generated with nucmer (MUMmer4 v4.0.1; --maxmatch -l 50 -b 500-c 200; (Marçais et al., 2018)) and filtered with delta-filter to retain alignments ≥200 bp at ≥90% sequence identity. Structural and sequence variations were identified with SyRI v1.6.3 (Goel et al., 2019). Intrachromosomal structural variants ≥20 kb were visualised with plotsr v1.1.1 (Goel and Schneeberger, 2022).

### Genome coverage plots and SNP density analysis

Filtered reads used in assembly (≥40 kb ≥Q20) and gap closure (≥100 kb ≥Q7) were independently mapped to both *E. recurva* haplotypes with minimap2 v2.29 (Li, 2018). Coverage was plotted using the Jvarkit v2024.08.25 tool WGSCoveragePlotter (Lindenbaum, 2015). For the high coverage ≥40 kb ≥Q20 subset, SNPs were called against haplotype 1 with longcallD v0.0.10 (Gao et al., 2026). Each chromosome was tiled into non-overlapping 10 kb windows and SNPs per window counted with BEDTools v2.31.1 (Quinlan and Hall, 2010) via pybedtools v0.12.0 (Dale et al., 2011), then plotted with matplotlib v3.10.8 (Hunter, 2007). Per-chromosome density was plotted as line graphs with counts capped at 1,000 SNPs per window (10% heterozygosity). The final genome-wide density was summarised as a single heatmap with colour saturating at 100 SNPs per window (1% heterozygosity).

## Data Availability

Raw ONT reads and genome assemblies have been deposited at the European Nucleotide Archive (ENA) accession [*****] (to be assigned upon submission). Genome assemblies, organelle sequences, and annotation files are available at [****URL].

## Acknowledgements

The authors acknowledge and thank Australia’s First Nations Peoples, the Traditional Custodians of the lands on which *Eucalyptus recurva* grows, and pays respect to Elders past, present, and emerging. The author thanks Genevieve Wright (NSW Department of Climate Change, Energy, the Environment and Water) for collection of plant material under the relevant scientific licence.

